# Cryptic cycling by electroactive bacterioplankton in Trout Bog Lake

**DOI:** 10.1101/2024.07.11.603116

**Authors:** Charles N. Olmsted, Mark Gahler, Eric Roden, Ben Peterson, James Lazarcik, Patricia Q. Tran, Maureen Berg, Donald A. Bryant, Danielle Goudeau, Rex R. Malmstrom, Mohan Qin, Katherine D. McMahon

**Author notes:** Corresponding author: (, 1550 Linden Dr, Madison, WI 53706). Data Availability: Sequencing data are published under the NCBI Bioproject ID PRJNA1018295. Metagenome assembled genomes are published at https://osf.io/kmj2s/. CRediT (Author Contributions): **Charles N. Olmsted:** Conceptualization (lead); Data Curation (lead); Formal Analysis (lead); Funding Acquisition (supporting); Investigation (lead); Methodology (lead); Software (lead); Visualization (lead); Validation (lead); Writing – Original Draft Preparation (lead); Writing – Review & Editing (lead). **Mark Gahler**: Conceptualization (supporting); Methodology (supporting); Software (supporting). **Ben Peterson**: Investigation (supporting); Formal Analysis (supporting); **Eric Roden**: Conceptualization (supporting); Formal Analysis (supporting); Validation (supporting); **James Lazarcik**: Formal Analysis (supporting); Methodology (supporting); Writing – Review & Editing (supporting). **Patricia Q. Tran**: Visualization (supporting), Writing – Review & Editing (supporting). **Maureen Berg**: Formal Analysis (supporting); Investigation (supporting); Methodology (supporting); Writing – Review & Editing (supporting). **Donald A. Bryant**: Investigation (supporting); Writing – Review & Editing (supporting). **Danielle Goudeau**: Formal Analysis (supporting); Investigation (supporting); Methodology (supporting). **Rex R. Malmstrom**: Investigation (supporting). **Mohan Qin**: Conceptualization (supporting); Validation (supporting). **Katherine D. McMahon**: Conceptualization (supporting); Funding Acquisition (lead); Resources (lead); Writing – Review & Editing (supporting).

## Abstract

The genetic potential for extracellular electron transfer (EET)-based metabolism has been shown to be a prevailing feature of humic lakes where bacterioplankton may be able to use EET to cycle dissolved organic matter (DOM) extracellularly between oxidized and reduced states, but measurable abiotic features resulting from this phenomenon have yet to be demonstrated. We observed an anoxygenic photosynthetic *Chlorobium* sp. bloom each summer in Trout Bog Lake in northern WI, USA. Given this bloom’s characteristics, we hypothesized that EET-based metabolisms of *Chlorobium* sp. and accompanying bacteria cycle DOM between oxidized and reduced states with seasonal or diel-timescale oscillations; therefore, we anticipated this could be measured by weekly and subdaily sampling. We collected vertical profiles on these timescales using a multiparameter sonde, including oxidation-reduction potential measurement, and we assayed for inorganic electron donors. We also developed and deployed a buoy to measure electric current flow between many pairs of electrodes simultaneously. Using metagenomics analyses, we examined the EET genes and other oxidoreductases of bacteria from water column samples at select depths and from biofilms that developed on electrodes at similar depths. Our results indicate the occurrence of diel electron cycling between phototrophic oxidation (electrotrophic metabolism) and anaerobic respiration (electrogenic metabolism), likely involving DOM. We also observed a gradual seasonal increase in hypolimnion oxidation-reduction potential. These diel and seasonal patterns have implications for carbon emissions and the ecology of electroactive bacteria in lakes.

**IMPORTANCE:** We investigated the physical, chemical, and redox characteristics of a bog lake and electrodes hung therein to test the hypothesis that dissolved organic matter is being cycled between oxidized and reduced states by electroactive bacterioplankton powered by phototrophy. To do so we performed field-based analyses on multiple timescales using both established and novel instrumentation. We paired these analyses with recently developed bioinformatics pipelines for metagenomics data to investigate genes that enable electroactive metabolism and accompanying metabolisms. Our results are consistent with our hypothesis and yet upend some of our other expectations. Our findings have implications for understanding greenhouse gas emissions from lakes, including electroactivity as an integral part of lake metabolism throughout more of the anoxic parts of lakes and for a longer portion of the summer than expected. Our results also give a sense of what electroactivity occurs at given depths and provide a strong basis for future studies.

## INTRODUCTION

Small inland lakes and especially their sediments are disproportionately important sources for greenhouse gas emissions and carbon burial (Dean and Gorham, 1998; Jonsson *et al*., 2001; Juutinen *et al*., 2003; Ask *et al*., 2012), and the overlying water column was recently proposed to control emissions in part through the metabolism of electroactive bacterioplankton (Olmsted *et al*., 2022). Extracellular Electron Transfer (EET), the process of routing electrons through the outermost membranes of cells, is considered prevalent in many anoxic and redox-transition environments where bacteria can use EET to perform redox reactions with colloidal or precipitated substances like iron, other metals, and quinone moieties in organic matter (Lovley *et al*., 1999; Coates *et al*., 2002; Kappler *et al*., 2004; Roden *et al*., 2010). The prevalence of EET in these systems can lead to measurable abiotic environmental features. As a premier example, banded iron formations in Precambrian marine basins prior to the rise of oxygen in Earth’s atmosphere are thought to be signatures of EET associated with anoxygenic phototrophic iron-oxidation and deposition (Hegler *et al*., 2008; Ward and Shih, 2022). Analogous features that one might expect to result from EET in lake water columns can be obscured by fast nutrient cycling rates called “cryptic cycling,” like that of iron and manganese within the anaerobic layer of the low-iron lake, Lake Cadagno (Berg *et al*., 2016). Besides metals, a prominent substrate for EET in many lakes may be extracellular Dissolved Organic Matter (DOM), considering that interfacial oxygen-mixing events can regenerate oxidized DOM for anaerobic respiring bacteria (Lau *et al*., 2017) and that high-DOM lakes are often quite enriched in bacteria with the genetic potential for EET (He *et al*., 2019; Olmsted *et al*., 2022).

This study focuses on Trout Bog Lake, a small, low-iron peat-bog lake with a characteristically high DOM content: roughly 20 mg C/L. Much of the DOM originates from the surrounding wetlands (Maizel *et al*., 2017), composed of variably sized recalcitrant lignin-derived and *Sphagnum* moss-exuded aromatic molecules (Walpen *et al*., 2018). These phenolic and quinoid molecules can lead to a relatively high electroactive capacity (electron accepting and donating capacity). Trout Bog Lake in particular has a high level of this aromaticity (Maizel *et al*., 2017) and electroactive capacity (He *et al*., 2019) that is likely comprised of usable EET substrates comparable to the common quinone-modeling compound anthraquinone-2,6-disulfonate (AQDS), which would benefit electroactive bacteria (Bai *et al*., 2020). Organisms that use reductive EET (redEET), for example to reduce AQDS, are called electro*gens*. In Trout Bog Lake, a *Geothrix* sp. was previously found to express multiple redEET genes (Olmsted *et al*., 2022), and *Geothrix* spp. are commonly reported electrogens that consume various simple organic acids and perform anaerobic respiration (Coates *et al*., 1999).

Electro*trophs* are capable of oxidizing extracellular substrates *via* oxidative EET (oxiEET). This may include iron oxidation as a source of reducing power for anoxygenic photosynthesis (Guzman *et al*., 2019; Peng *et al*., 2019), potentially supporting heterotrophs by providing carbon, energy, and redox equivalents (*e.g*., Liu *et al*., 2013). The two distinct populations of anoxygenic phototrophic *Chlorobium* spp. in Trout Bog Lake (Berg *et al*., 2021) are both represented by metagenome assembled genomes (MAGs) that harbor three homologues of Cyc2, an iron-oxidizing EET protein (Keffer *et al*., 2021). These *Chlorobium* spp. were also found to transcribe genes for at least one Cyc2 homologue far more than their oxidoreductases for other electron donors like hydrogen or sulfur compounds (Olmsted *et al*., 2022). Only one *Chlorobium sp.*, referred to by Berg *et al*, 2021 as “GSB-B” which corresponds to “3m_metabat_bin_57” (Tab. S1), blooms over the course of the summer just below the oxycline. Regarding nocturnal metabolism, there exists some evidence that members of *Chlorobium* can ferment (Badalamenti *et al*., 2014) and less evidence for respiration (Badalamenti *et al*., 2013). The *Chlorobium* spp. in Trout Bog Lake have genes that suggest fermentation is possible, (*e.g.*, degrading glycogen stores), but also have the genes for nitric oxide reduction, oxygen reduction, and even 2–3 genes that match membrane proteins—FmnB+DmkB or FmnA+DmkAB+Ndh2 depending on the MAG—that catalyze electron transfer into the quinone pool in other organisms to selectively use particular electron acceptors (Unden and Bongaerts, 1997; Light *et al*., 2018). Although it is uncertain as to whether Trout Bog *Chlorobium* spp. are actively metabolizing at night, we understand that during the day they require a source of reducing power, *i.e.* electron intake, to reduce CO2 to biomass for anoxygenic photosynthesis.

Considering the electroactive bacteria and environmental characteristics of Trout Bog Lake, we hypothesized that anoxygenic photoautotrophs, like *Chlorobium* sp., generate zones of oxidized DOM *via* electrotrophy that can be restored by respiring electrogens like *Geothrix* sp. Also, EET can be detected in real time because of the resulting electrical features (*e.g.* Rowe *et al*., 2021; Upadhyay *et al*., 2021). We therefore suspected we might be able to observe this activity either seasonally or on a diel scale in Trout Bog Lake due to the high proportion of the *Chlorobium* sp. that establishes by late summer in the redoxcline. We combined weekly seasonal measurements, automated electrical measurements, chemical assays, and metagenomics with intensive diel profiling to investigate electron cycling between anoxygenic phototrophic electrotrophy and electrogenesis with respect to the microbial ecology and DOM of Trout Bog Lake.

## EXPERIMENTAL PROCEDURES

### Trout Bog Lake Characteristics

Trout Bog Lake is a darkly stained bog lake on the northwest periphery of Trout Lake, Boulder Junction, WI and is part of the North Temperate Lakes Long-Term Ecological Research (NTL-LTER) program. This humic lake is located at the coordinates 46°02’28.2“N 89°41’10.6”W, has a surface area of 1.1 hectares, and has a maximum depth of 7.9 m. The warmer, oxygenated epilimnion is typically only represented by the top 1–2 m throughout the summer. In full sun, only 0.008% of photosynthetically active radiation could be detected at 2 m deep, 0% by 3 m (Fig. S0). As a bog lake, its edges are covered by floating *Sphagnum* sp. moss, *Gaultheria procumbens* wintergreen, and other typical bog flora, but a small portion of the boggy edge is not far away from solid forest floor under a mix of deciduous and coniferous trees.

### Metagenomics Collection and Extraction

We used all Trout Bog Lake MAGs from Olmsted *et al*., 2022 as well as additional MAGs (Tab. S1) generated (Text S2) from water and electrode biofilm samples (Tab. S2), for a total of 1301 MAGs—597 of which were identified as bacterial genomes over 50% complete and less than 10% contaminated. Both integrated and depth-discrete water samples were collected using a peristaltic pump into an opaque Nalgene sample container and transported in a cooler with ice packs to the University of Wisconsin Trout Lake Station. Water samples were filtered (0.2 µm polyethersulfone, Pall Corporation), frozen at –20 °C, and then transferred later to –80 °C, except for all 2021 water samples and 2020 electrode samples which were frozen directly at –80 °C. Phenol chloroform DNA extractions were performed on filters and electrodes as described (Peterson *et al*., 2020).

### Genome Analysis

Average percent identity (ANI) of MAGs was calculated using FastANI v1.3 (Jain *et al*., 2018). All MAGs were clustered under representative metagenomic operational taxonomic units (mOTUs) (Tab. S1) by using an identity cutoff of 98% and selecting the most complete and least contaminated MAG to represent each cluster. The displayed taxonomic abundance was derived from the reads of each sample realigned to mOTU representative sequences. Oxidative, reductive, and putative EET genes were determined using FEET (Olmsted *et al*., 2022), and the FEET pipeline definitions of redEET verses oxiEET are largely based on FeGenie (Garber *et al*., 2020). We used the METABOLIC V3 software (Zhou *et al*., 2022) to determine the genetic potential for various oxidoreductases and other metabolisms on MAGs over 50% complete and less than 10% contaminated, and we only considered MAGs of the same quality when summarizing gene counts in a given sample.

### Microghost Buoy Design and Deployment

We designed and deployed a buoy that automatically measured electrical current flow between pairs of unpoised electrodes. The circuit board design (File S1 MicroghostBoard.pcb), parts list, build information, and silly reasons for the name “Microghost” are available in Supplementary Text S1. We refer to the pairs of connected electrodes as “channels.” Prior to deployment, 16 pairs of 18 cm^2^ plain carbon cloth (1071 HCB, Fuel Cell Store, AvCarb) electrodes were weaved over 3 cm of exposed bundled (19 strands of 27 gauge) copper wire amounting to a surface area of 6.46 cm^2^ per electrode. Two buoy deployments occurred from July 21^st^, 2019 to August 13^th^, 2019, and the other from August 16^th^, 2020 to September 9^th^, 2020. A HOBO (Onset MX2202) datalogger device was secured on top of each buoy to measure sunlight intensity. Dissolved oxygen (DO) and temperature readings are from separate buoys placed roughly 10 m away using a miniDOT Logger (Precision Measurement Engineering) (Fig. S1D–F). Some 2020 buoy electrodes (Fig. S1A–C) were covered in calcium-alginate with pores with estimated diameters of 12−16 nm (Sergeeva *et al*., 2019). To coat electrodes, each was soaked in 1% sodium alginate, placed into 100 mM calcium chloride to cure for one hour, and rinsed with DI water. The same steps were repeated for a double layer. These electrodes were stored in 50 mM calcium chloride at 4 °C but rinsed in deionized water before deployment.

### Chemical and Biological Profiling

All chemical and biological samples were collected *via* peristaltic pump and later analyzed as detailed (Text S2). Depth-discrete cytometric counts of *Chlorobium* spp. and total cell counts were generated in 2018 by flow cytometry as described (Berg *et al*., 2021), but now we publish cell counts rather than only percent of total cells. Turbidity, pH, chlorophyll, temperature, and DO in 2017 (Fig. S2) were collected using an Exo1 multiparameter sonde (Yellow Springs Instruments). In 2018 and beyond, the same parameters—except oxidation-reduction potential (ORP) instead of chlorophyll—were measured using a ProDSS multiparameter sonde (Yellow Springs Instruments). Our diel ProDSS vertical profiling procedure was as follows, all taken by a single researcher to minimize human-derived differences between profiles. All parameters were calibrated once just prior to the sampling trip. The sonde was set outside the water for 1 min and then given 5 min to equilibrate in the water at a depth of zero meters before dropping the sonde at constant rate of roughly 1 cm every 1.5 s, logging values every 2 s. Each vertical profile took about 15 min, and thus the beginning measurement for each replicate profile was about 21 min apart. Values were logged only as the sonde was lowered.

## RESULTS

### Sunlight-driven oxidation where anoxygenic phototroph *Chlorobium* sp. is abundant

We performed subdaily multiparameter vertical profiling in 2021 on Trout Bog Lake to test whether there are observable oscillations on a diel timescale in oxidation-reduction potential (ORP) and other factors within the *Chlorobium* sp. bloom. At 2.0 m deep we observed significant diel shifting of the ORP (Fig. 1B,C), a measure of the potential between the electroactive components of the water interacting with a platinum electrode and a Ag/AgCl reference electrode on the ProDSS sonde. In the region with the most diel variation of ORP, 1.0–2.5 m deep, ORP decreased gradually at night, reaching its lowest potential just before dawn.

**Figure 1.**
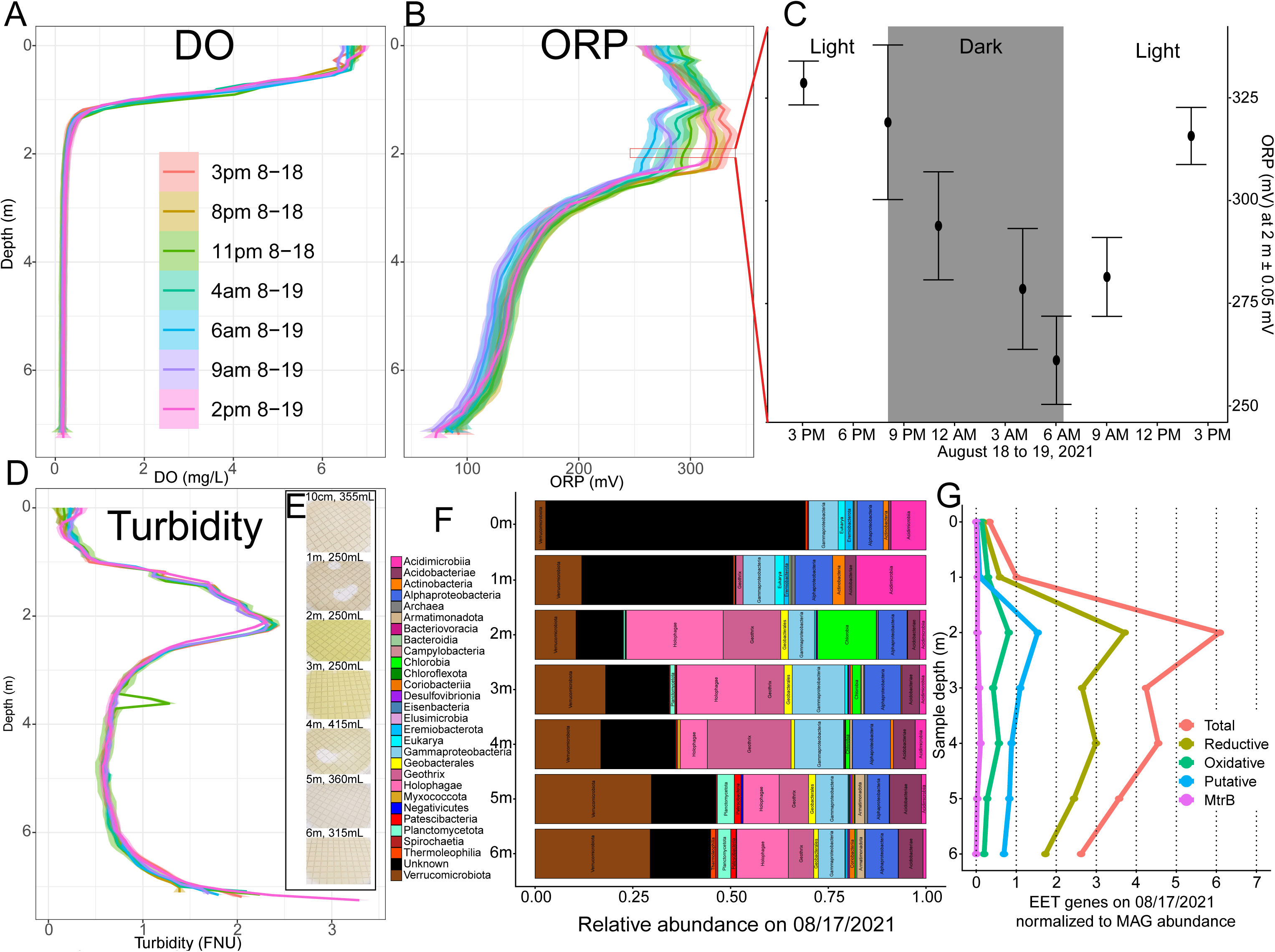
Observance of light-driven oxidation during diel sampling campaign on Trout Bog Lake including vertical profiles for (A) dissolved oxygen (DO), (B) oxidation reduction potential (ORP) and (C) significant ORP change at 2 m deep, (D) turbidity, (E) bacterial filters and (F) resulting relative abundances of MAGs, and (G) counts of EET genes normalized to each MAG’s relative abundance. Profiles were collected by slowly descending a ProDSS sonde, each timepoint is an average of three profiles, each 20 min apart. Bacterial filters were pictured by a phone camera just prior to freezing at -80°C, and later subjected to extraction and metagenomics. Normalized EET gene counts were calculated by multiplying each MAG’s number of EET genes by the relative abundance of that MAG and adding all such values together per sample. Measurements and materials were collected between August 17th, 2021, and August 19th, 2021.

After only 2–3 h of daylight, an obvious increase in ORP (oxidation occurring) initiated at 2.0 m, the depth the genus *Chlorobium* was found to be most abundant in 2021 (Fig. 1E,F). Later in the day, the ORP of the entire region with diel variation returned to more oxidized values as observed prior to dusk the previous day (Fig. 1B,C).

### Unexpected seasonal oxidation in Trout Bog Lake hypolimnion

We collected vertical profiles of chemical and physical data using a ProDSS multimeter sonde over several summers (Fig. 2, Fig. S3), and we assayed for chemicals including sulfide, phosphorus, manganese, iron, nitrate, and ammonium at various depths (Tab. S3, Fig. S4). Since ORP is a value that depends on the ratios and electroactive tendencies of reductants and oxidants, including DOM (Struyk and Sposito, 2001; Aeschbacher *et al*., 2011; Kane *et al*., 2019), we use ORP as a proxy for the redox state of DOM. We make this assumption because electroactive capacity of DOM in Trout Bog Lake (He *et al*., 2019) is at least one order of magnitude higher than each other measured chemical that likely also contributes to the ORP, such as iron and sulfide (Tab. S3, Fig. S4). The expectation is that in a strongly stratified, anoxic body of water with diverse anaerobic respiring bacteria, substances in the water will become more and more reduced as oxidized substrates are respired (reduced) roughly in sequential order (Avetisyan *et al*., 2019) down the redox ladder (Chen *et al*., 2017), leading to a gradually decreasing ORP from the onset of spring stratification until fall mixing. However, we observed that the ORP of the hypolimnion, including the anoxic water down to a meter above the sediment, achieved its most reduced state within weeks after spring mixing and thereafter trended towards becoming more oxidized. We observed this for the summers of 2018, 2019, and 2021 (Fig. 2D–F). Summer of 2020 had too few sample dates to detect trends, due to pandemic-imposed fieldwork restrictions. From our earliest spring profile to the latest in August 2018, 2019, and 2021 the average ORP from 2–6 m respectively increased 166 mV, 110 mV, and 128 mV.

**Figure 2.**
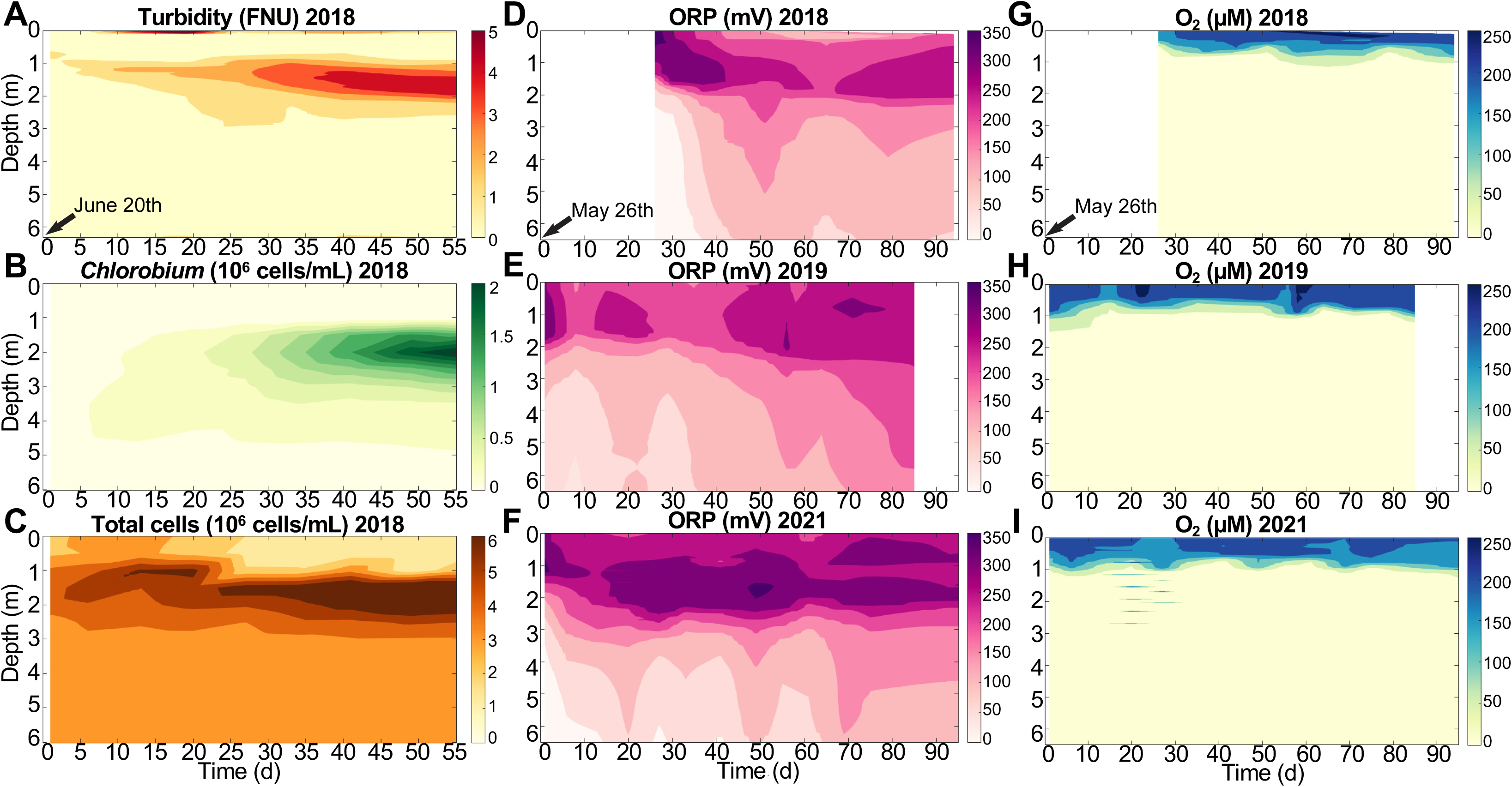
Seasonal measurements from Trout Bog Lake including (A) turbidity, (B) cytometric counts of *Chlorobium* and (C) total cells in 2018, and (D3F) oxidation reduction potential (ORP), and (G3I) dissolved oxygen for 2018, 2019 and 2021 summers. The beginning date for each column of graphs is indicated, and blank space is added so date for each year is aligned vertically. Cytometric data was collected by peristaltic pump. All other data was collected by slowly dropping down a ProDSS sonde.

### Comparison of water column and electrode biofilm microbial communities

We collected cytometric data (Berg *et al*., 2021) and generated metagenomics data from water samples collected as part of the NTL-LTER time series and from additional depth-discrete samples collected in 2021 (Fig. 1E, Tab. S2). This sampling yielded MAGs similar to those found by previous depth-integrated sampling of Trout Bog Lake (Olmsted *et al*., 2022) (Fig. 1F, Tab. S1). Except, at 0–1 m deep, most sequences (51% and 23%, respectively) mapped to a 225.5 kb single-contig MAG, 0m_metabat_bin_56, that is likely a virus (Fig. S11) in the order Caudoviricetes, based on the MVP pipeline (gitlab.com/ccoclet/mvp). Otherwise, the most common bacteria were in clades expected to respire oxygen and consume organic matter, such as Actinobacteria and *Novosphingobium*. However, one of the more abundant epilimnion-dwelling bacteria was classified within *Steroidobacteraceae*, and it harbored both oxiEET and redEET genes—that is, respectively, genes encoding proteins expected either to oxidize or reduce extracellular substances through EET. The most abundant bacterial family found at 2 m and below was *Holophagaceae*, the most common genus of which overall was the electrogen *Geothrix*, and each harbored 7–10 redEET genes. The next most abundant bacterial genus was *Chlorobium*, found maximally at 2 m. Another high-abundance genus was *Polynucleobacter*, a *Burkholderiaceae* common in lakes (Newton *et al*., 2011), and 75% of these MAGs had oxiEET genes. Trout Bog also had a high proportion, increasing by depth, of *Verrucomicrobiaceae*, which are thought to consume (poly)saccharides primarily, and 85% of such MAGs had two or more EET genes which is consistent with past studies (He *et al*., 2017).

We compared the number of EET genes between MAGs found in water samples from 2019, 2020, and 2021 (Fig. 1, Fig. S5) with MAGs found on four of the electrodes from the 2020 Microghost Buoy (described in more detail below) (Fig. 3). The electrode biofilms were enriched with bacteria containing EET genes. Specifically, the number of EET genes normalized by the relative abundance of each MAG was higher on electrodes than in the water, except for the electrode deployed at 2.25 m which had a far lower abundance-normalized EET gene count than the water at 2 m (Fig. 3B). On the electrodes overall, more oxiEET than redEET genes were recovered from the cathode in the epilimnion while more redEET than oxiEET genes were recovered from anodes at 5.5 m in the hypolimnion water, by both metrics of diversity (Tab. S1) and relative abundance (Fig. 3B). *Geobacter* sp., an electrogen with numerous redEET genes, was also enriched on anodes at 5.5 m compared to surrounding water (Fig. 1F, Fig. 3A). Most of the diversity in electrode biofilms consisted of electrode microbes; that is, any MAG for which its mOTU (98% ANI) was recovered at the highest quality from that or another electrode.

**Figure 3.**
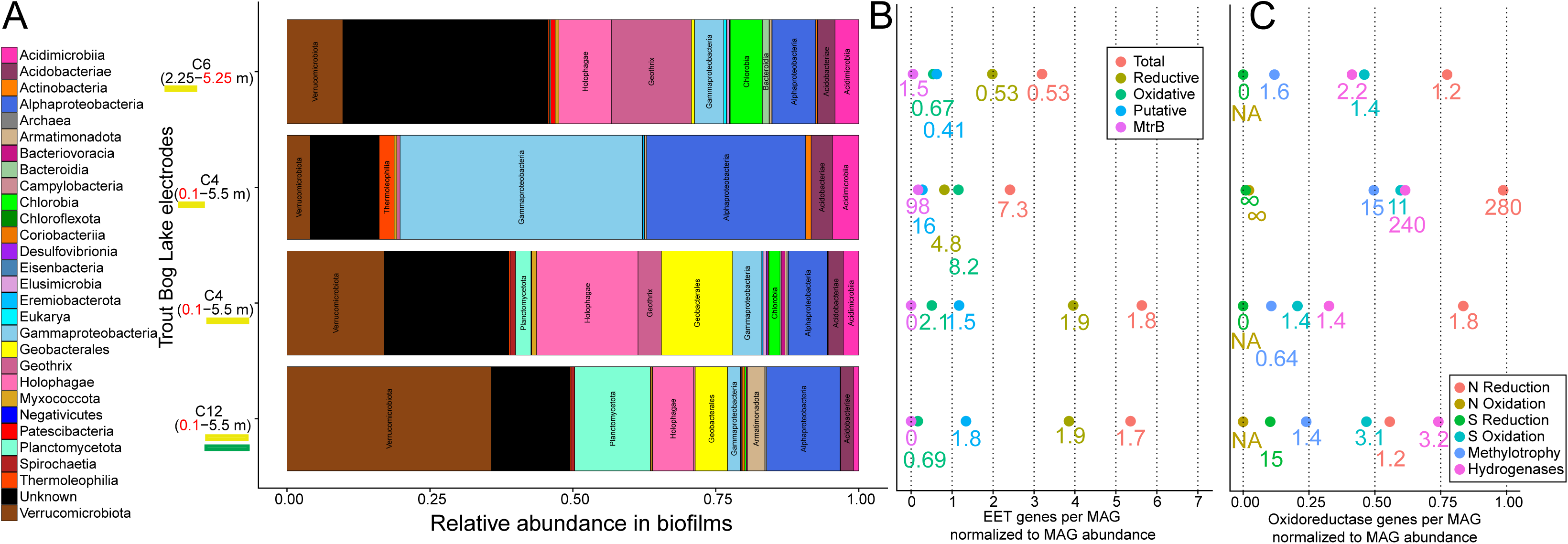
Microghost Buoy electrode biofilm MAG (A) taxonomy, (B) EET genes, and (C) other oxidoreductase genes normalized to the relative abundance of MAGs. Each row represents data from one electrode in a pair of electrodes connected by 4.75 Ω, a “channel,” and the depth of the represented electrode has a yellow underline. The green underline indicates the electrode covered in porous calcium alginate. The channel number above the anode (black text) and cathode (red text) depths corresponds to the 2020 channel numbers in Supplementary Figure S1. Normalized gene counts were calculated by multiplying each MAG’s number of the given gene type by the relative abundance of that MAG and adding all such values together per sample. Relative abundance represents the average metagenomic read mapping for all mOTUs in the given clade, and displayed subclades were not double counted in higher clades. The point labels represent the point’s value divided by the value found for water samples near the same depth, 2 m water sample for the 2.25 m electrode and the average of 5 m and 6 m water samples for the 5.5 m-deep electrode. NA represents when the value was zero for both while infinity symbols are for when only the water sample value is zero.

However, the most dominant electrogen of the water column, *Geothrix* sp., was an electrode microbe by that definition, and dominant members of the *Chlorobium*-zone, particularly *Chlorobium* sp., *Geobacter* sp., and *Geothrix* sp, and *Holophaga* spp., were among the most dominant members in anodic biofilms.

### Influence of sunlight on electrical current between *in situ* electrode pairs

To get a general sense of how the interactions between electroactive bacterioplankton, water chemistry, and a conductive surface might vary day to day, we deployed an automated buoy (Fig. 4) that measured the microamperage between pairs of unpoised carbon cloth electrodes connected by a low (4.75 Ω) resistance. We deployed this buoy alongside other automated buoys measuring physical variables. There was a general increase in electron flow over the course of the time series (Fig. S6, Fig. S7) which is consistent with the growth of EET-capable bacterial biofilms. For channels with a deep (5±1 m) electrode in anaerobic water and a shallow electrode in oxygen-rich water, electrical current was as expected: overall positive as electrons flowed from the deep anode towards the shallow cathode. A more detailed explanation of observed electrical patterns is provided (Text S2), but generally sunlight intensity correlated to decreased electron transfer from all anodes located in *Chlorobium*-abundant depths (2.5±0.5 m) to corresponding cathodes more so than for the channels with deeper anodes.

**Figure 4.**
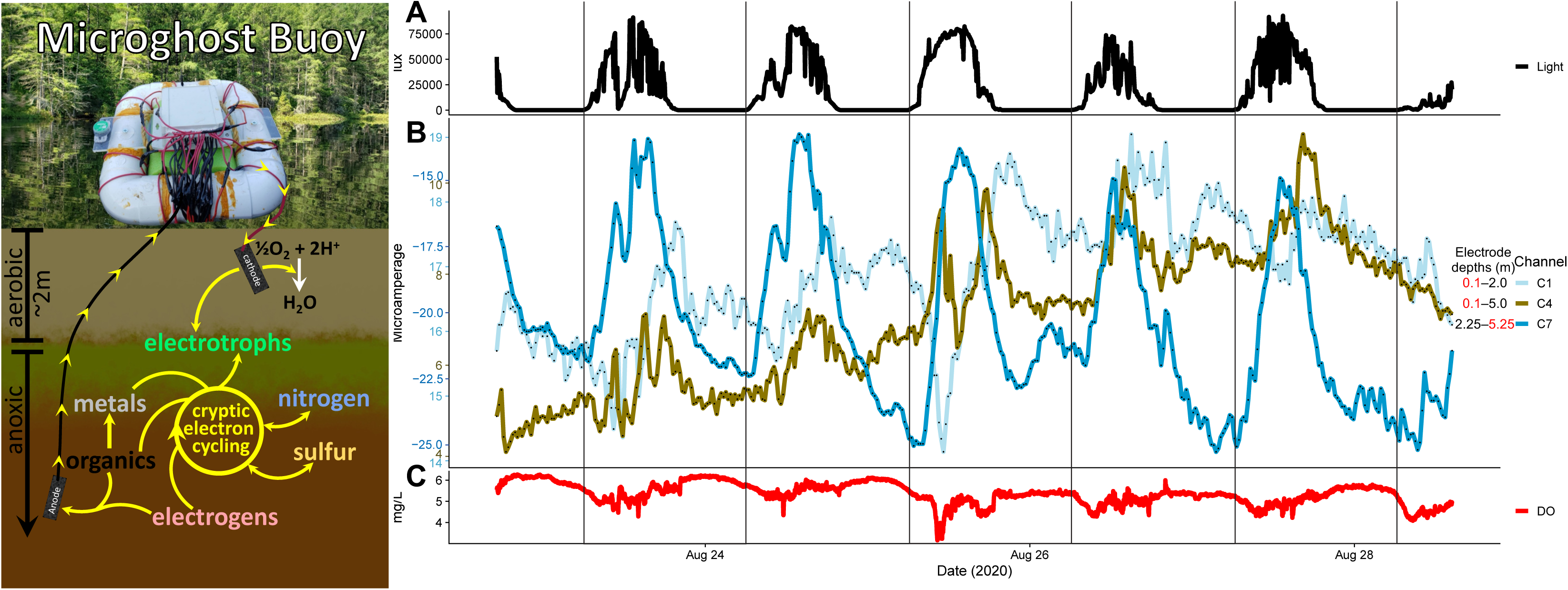
Concept figure of Microghost Buoy function (Left) and timeseries data from 2020 buoy deployment including (A) surface light intensity, (B) electrical current measurements for all channels, (C) dissolved oxygen (DO) measured at 20 cm deep. Direction of electron transfer across 4.75 « is indicated by yellow arrows. The background picture is of Trout Bog Lake, and a picture of the Microghost Buoy during its 2020 deployment is pasted on top. Depth (m) of cathodes (red text) and anodes (black text) is as indicated. All electrodes were plain carbon cloth connected by insulated copper wire. For all electrodes deeper than 10 cm, the wires were bundled together, and the electrodes were splayed out horizontally from the main bundle as to not risk touching each other. All 10 cm-deep cathodes were fixed in place under the buoy9s shadow.

To test for diel variations in the *Chlorobium*-abundant zone absent the impacts of near-surface dynamics, we connected some channels (Fig. S1A,B,C6,C7,C10) from 2.5±0.5 m to 5±1 m, where we did not expect as much diel variation. These channels (including Channel 7 in Figure 4B) showed negative currents because the electrodes that acted as the anode or cathode were opposite of what we anticipated—*i.e.* the direction of electron flow was from 2.5±0.5 m towards 5±1 m. For these channels, unlike other channels, positive correlations regarding the diel component of current indicate decreased electron flow away from 2.5±0.5 m.

We also performed multivariate linear regression modeling with the dependent variables of surface light, dissolved oxygen (DO) at 20 cm, and/or temperature at 20 cm against the independent variable, the diel component of current, for the three channels in Figure 4. The amperages are displayed for the same period used for this modeling, the middle third of the time series (Tab. S4). For all three channels during the displayed time, all three dependent variables were significantly correlated. However, light was the best predictor of, and negatively correlated to, electron flow from the anodic electrodes of Channels 1 and 7 that were positioned within the *Chlorobium*-enriched zone. This was unlike the 5 m-deep anode of Channel 4 for which light was positively correlated. Temperature was next best for Channels 1 and 7, adding some predictive power to the models, but it was the best predictor for Channel 4. Temperature positively correlated to electron flow from anodes for all three channels. The least predictive of the three variables, DO, was positively correlated to electron flow from the anodes of Channels 1 and 7, negatively for Channel 4. We recognize the dependent variables covaried with each other over this modeling window, except temperature and DO at 20 cm (Fig. S1D1,E1,F1).

## DISCUSSION

### Cryptic electron cycling by anoxygenic phototrophs and electrogens

The EET gene enrichment in bacteria at *Chlorobium*-abundant depths (Fig. 1E–G), correlations in electrical current production with sunlight, and, most convincingly, the diel fluctuation in ORP all provide evidence for EET-based electron cycling between *Chlorobium* sp. and electrogens in the upper layers of the Trout Bog Lake hypolimnion. In fact, the mere maintenance of such high ORP below the oxycline and in the domain of reductive heterotrophs (Lau *et al*., 2017) provides an obvious clue that an abundant organism (*e.g., Chlorobium* sp.) was performing oxiEET (Fig. 2). Given the abundance of DOM as an electroactive species in Trout Bog water (He *et al*., 2019, Tab. S3), it is likely that the oxidation of reduced quinone moieties on DOM played a key role in this observed oxiEET activity. We have not yet determined *via* assays that would require isolated *Chlorobium* sp. whether DOM is a direct or an indirect electron donor to the *Chlorobium* sp. oxiEET proteins. However, reduced DOM (*e.g.* AQDS) can quickly reduce oxidized iron (Kappler *et al*., 2004; Bai *et al*., 2020); thus, even if ferrous iron is the dominant substrate transferring electrons to the oxiEET proteins, coupled redox cycling of DOM and iron could sustain and amplify *in situ* cryptic electron cycling between electrotrophic *Chlorobium* sp. and electrogenic organisms such as *Geothrix* sp. Considering this, while taking into account potential redox chemistry (Tab. S3, Fig. S8) with the platinum ORP electrode (Struyk and Sposito, 2001; Aeschbacher *et al*., 2011; Kane *et al*., 2019), the net redox difference recorded by ORP (Fig. 1B) likely does not fully capture the gross rate of electron flux linked to *Chlorobium* sp. metabolism. This is consistent with the relatively high cooccurrence of oxiEET and redEET gene expression (Olmsted *et al*., 2022) as well as photosynthesis and respiration gene expression (Linz *et al*., 2020) in Trout Bog Lake.

The fact that both oxiEET and redEET genes were enriched in genomes and in the community at 2 m supports the occurrence of EET-driven electron cycling in the hypolimnion. Based on the average number of EET genes per MAG normalized by MAG relative abundance (Fig. 1G), the potential for EET was greatest at 2 m and decreased with depth in the hypolimnion. Without normalizing by abundance, EET genes per MAG increased below 2 m (Fig. S9A). This supports our conceptual model describing how EET operates in Trout Bog Lake: at the intersection of the light and anoxia gradients, anoxygenic phototrophic electrotrophs, particularly *Chlorobium* sp., proliferate alongside electrogens like *Geothrix* sp., providing both primary productivity-derived organic compounds, oxidized iron, and presumably oxidized DOM (Fig 4, Fig. S10). As these extracellular electron acceptors (*e.g.*, oxidized quinone moieties) and labile organic compounds become scarcer deeper in the water (further away from *Chlorobium* sp.), bacteria that rely on EET-independent metabolisms (Fig. S9D) become more prevalent.

Meanwhile, bacteria that still rely on EET may require more EET gene redundancy and/or expression to achieve a higher concentration and diversity of EET membrane proteins to access the increasingly scarce yet likely diverse EET substrates.

The diel current waveforms of Microghost Buoy channels in conjunction with the overall increased current flow over time as biofilms grew (Fig. S6, S7) may relate to the cryptic cycling of electroactive substrates and EET metabolism by acting as a sensor for the diel activity of *Chlorobium* sp. We expected that if we connected an electrode in anoxic, reduced bog water to another electrode near the surface in more oxidized, oxygenated water, electrons would flow upwards as both reduced DOM and electrogens in the anoxic region donate electrons to the electrode (Fig. 4), similar to the microbial snorkel concept (Matturro *et al*., 2017). Overall, this was the case, even for channels connected from the oxygenated water to the even higher-ORP yet anoxic *Chlorobium*-abundant depths (∼2 m). This result suggests that the electrode biofilms contributed more to current generation than reduced molecules in the surrounding water interacting directly with the electrode surface.

Several factors clearly influenced electrical currents, but the decreases in total electron donation to electrodes from biofilms in the *Chlorobium*-abundant zone were most strongly determined by sunlight and, as we suggest, *Chlorobium* sp. “photoelectrotrophy.” This is exemplified by our multivariate analysis and how channels’ electrical patterns follow increases and decreases in sunlight (Fig. 4). We note how the current of Channel 1, which had an aerobic cathode subject to surface effects and an anode in the range of *Chlorobium* sp., decreases shortly after sunlight becomes available for photoelectrotrophy. This pattern is even more apparent for Channel 7, which was affected only by happenings within the range of *Chlorobium* sp. and below: whenever sunlight increased, within minutes, the negative current began shifting towards zero, and whenever clouds obscured the sun, the negative current spiked as electrons otherwise occupied by photoelectrotrophy were permitted to donate to the anode. Meanwhile, the fluctuations of Channel 4 likely represented electrochemistry on its aerobic cathode, considering the expectedly steady donations from electrogens and reductive chemistry to its deep anode (Fig. 4). The possibility of electroactivity on aerobic cathodes was considered, particularly as biofilms grew, likely providing microenvironments for both anaerobic electrogens and microaerophilic electrotrophs. Light-dependent electrogenesis has been shown in some organisms (Li *et al*., 2015) even including some cyanobacteria to compensate overcharged photosystems (Pisciotta *et al*., 2011), and this may have impacted some of the electrical measurements. This idea is also supported by strong negative correlations with sunlight for cathodes at 70 cm where one expects more aerobic phototrophs based on 2017 chlorophyll data (Fig. S2).

The consistently repeatable phenomenon of downward current flow from evidently anodic electrodes in *Chlorobium*-abundant depths (2.5±0.5 m) to unexpectedly cathodic electrodes at more reduced, deeper depths (5±1 m) suggests there was something—whether biotic or abiotic—around 5±1 m and/or on the electrodes in that depth range capable of accepting electrons at the prevailing redox potential maintained by the shallow electrode (Fig. 4B, Fig. S1A,B,C6,C7,C10). Unfortunately, the electrode from which we attempted to sequence biofilm DNA to see if these deep cathodes were enriched for electrotrophs, yielded abnormal traces and thus had to be excluded from shared-lane sequencing. However, our available pelagic metagenomes yielded potential explanations. The greater abundance of electrogens at 2 m, indicated by increased turbidity and MAG relative abundance, explains why their electrogenic respiration might overpower the electrogenic respiration at deeper depths. Furthermore, the electron acceptance from electrodes at 5±1 m could be due to electrotrophic chemoautotrophy, consistent with the uptick of oxiEET at 4 m (Fig. 1G, Fig. S9A) from organisms in the order Burkholderiales (particularly *Ferrovaceae* and *Rhodoferax*) and in the families *Magnetospirillaceae* and *Acetobacteraceae* (Tab. S1). Given the potential for chemoautotrophy in these organisms, the substrate availability at 5±1 m, and supported in part by differing abundances of hydrogenases between EET+ and EET– organisms (Fig. S9C,D), the terminal electron acceptor seems likely to be CO2. The presence of sulfur and nitrogen-compound oxidoreductases (Fig. S9B) suggest these substrates may also be used, but sulfate and especially nitrate are not—nor expected to be—*statically* abundant. However, dynamic abundance is conceivable given cryptic cycling.

### Influence of electrotrophic *Chlorobium* spp. oxidation on nitrogen and sulfur cryptic cycles

Diverse sulfur and nitrogen-cycling microbes capable of EET that may be sensitive to changes in the overall redox state would be influenced by the oxidative activity of *Chlorobium* spp. 2 m and below. *Chlorobium* spp. may also influence them more directly through ecological interactions like competing for reduced substrates or providing reducible and consumable substrates. Notably, we observed slight yet significant diel variability in ORP between 3.5 to 6 m (Fig. 1B), and, while DOM is likely the dominant electroactive component, ORP is a summary measurement of many electroactive substrates including sulfur, nitrogen, and other compounds (Fig. S4). There may be multiple explanations for why we observed diel variability in ORP so deep in the bog, such as minor levels of *Chlorobium* spp. oxiEET and/or sulfur oxidation. There is a low yet measurable count of *Chlorobium* spp. in this range (Fig. 2B), potentially photosynthesizing in very dim light (Parkin and Brock, 1980) considering our photometer may not have been sensitive enough to detect trace scattered red and infrared light (Fig. S0).

Otherwise, the shift in ORP at 3.5 to 6 m could even conceivably have resulted from interactions with phototrophic processes from the day prior. Closer to the sediment, sulfide contributes more to decrease ORP, but there was a slight peak of sulfide at 3m (Fig. S4B). This suggests there may have been sulfur cycling, potentially by *Chlorobium* spp. and other organisms, and the concurrence of oxidoreductases and EET genes raises the possibility that EET could have facilitated cryptic electron cycling between various redox states of sulfur, nitrogen (*e.g.* Coates *et al*., 2002; Li *et al*., 2020), iron, and DOM. The differences in depths where one finds more EET+ verses EET– organisms with sulfur and nitrogen oxidoreductases (Fig. S9) also supports this possibility. Based on abundance at 4m and their genes, multiple bacteria that might play a role this EET-involved cryptic redox cycling were identified. Metagenomics suggests *Ferrovaceae* may be capable of oxiEET, redEET, methanol oxidation, carbon fixation, sulfide oxidation, and thiosulfate redox. *Terracidiphilus* sp. seems capable of oxiEET, redEET, hydrogen metabolism, sulfur(ide) oxidation, nitrate(ite) reduction, and acetogenic fermentation. *Magnetospirillaceae* appear capable of oxiEET, redEET, sulfide oxidation, and thiosulfate redox, methanol oxidation, and fermentation. Some of these organisms may also use oxiEET for their electron source while respiring fermentation products for an energy source or use oxiEET for chemoautotrophy.

### Oxidative EET controls summertime ecology in Trout Bog Lake, obstructing methanogens

Competition for simple organic acids resulting from the use of oxidized DOM as an electron acceptor for anaerobic respiration has been suggested to be a controlling factor for methanogenesis in humic-rich systems (Heitmann *et al*., 2007; Lau *et al*., 2017; Walpen *et al*., 2018). Lau *et al*. (2017) outlined this interaction regarding oxygen-linked redox cycling of DOM with anaerobic respiration near the oxycline. Our results indicate that photoelectrotroph-linked DOM oxidation could promote such respiration entirely within the anoxic hypolimnion. By profiling on a weekly scale throughout three summer seasons, we discovered that the ORP of Trout Bog Lake’s hypolimnion gradually became oxidized throughout our sampling period that ended in August. Considering the diversity of putatively electrotrophic bacteria that could have been responsible for this (Olmsted *et al*., 2022) (Tab. S1), we do not suspect the hypolimnetic oxidation was entirely due to *Chlorobium* spp., especially considering that the seasonal increase of ORP extends well below where *Chlorobium* spp. were most abundant (Fig. 1. Fig. 2).

Regardless of which suite of electrotrophic organisms was responsible, the key point is that the balance between oxidative and reductive EET-driven electron cycling shifted over the summer. In turn, the resulting increase in abundance of oxidized compounds as electron acceptors for anaerobic respiration would be expected to suppress methanogenesis.

Another way that hypolimnetic EET-driven electrotrophy could suppress water column methanogenesis is by simply providing a large quantity of anoxygenic photosynthetic and dependent electrogenic planktonic biomass with a means to remain alive until fall mixing.

Particularly in the case of a partial fall mix as observed at least sometimes (if not usually) in Trout Bog (Gorsky *et al*., 2021), all the living biomass which does not settle out beforehand will be exposed to oxygen, permitting more zooplankton grazing (Arvola *et al*., 1992), aerobic respiration, and oxygen-requiring degradative pathways. This may also be true for a full mix, but most of the highly fermentable biomass produced by primary production, is potentially above to just below the turbidity peak (Fig. 2) and exposure to oxygen-dependent mineralization might make some deeper carbohydrates fermentable (Leschine, 1995). Hypothetically, in the absence of either cryptic EET cycling or the overall hypolimnetic oxidation, the settling biomass from primary production would be not only richer in easily fermentable substrates but also on a more direct route—*i.e.* not suspended all summer long—to the zones occupied by methanogens thereby providing fermenters and *ipso facto* colocalized methanogens with a steady, labile feedstock.

## CONCLUSION

By combining seasonal and diel profiling with depth-discrete metagenomics and flow cytometry, previous metatranscriptomics, and automated electrical and physiochemical measurements in a darkly stained bog lake, we have provided evidence that anoxygenic photoautotrophs in such lakes can generate volumes of oxidized DOM *via* electrotrophy that can be re-respired by electrogens. This cryptic cycling and likely additional oxiEET may have the potential to cause gradual oxidation of DOM at the seasonal scale throughout nearly the entire hypolimnion. Considering the broad distribution of the blooming *Chlorobium* sp. and global prevalence of anoxygenic phototrophs carrying oxiEET genes (Garcia *et al*., 2021; Olmsted *et al*., 2022), our extended knowledge of this phenomenon yields widespread implications for ecosystem-scale methane metabolism and bacterial ecology.

We observed very clear diurnal oxidation in response to light initiating from the depth at which the dominant phototrophic electrotroph (*Chlorobium* sp.) in Trout Bog Lake is most abundant. Comparing the concentration and electroactive capacity of DOM to that of inorganic chemicals in Trout Bog Lake implies electroactive DOM contributes in a crucial way to the metabolism of oxiEET-capable anoxygenic phototrophs and dependent electrogens.

Our findings imply that even though organic matter is the main carbon and energy source that eventually feeds methanogenic activity, higher proportions of electroactive DOM, such as quinones produced by *Sphagnum* or microbially derived mediators, may produce unfavorable conditions for methanogens in anoxic hypolimnia because electroactive DOM is a regenerable substrate powering their competition. Electroactive DOM is not just regenerable by oxygen and mixing events, but it may also be re-oxidized by diverse anaerobic microbes, in particular *Chlorobium* spp. and other anoxygenic phototrophs. Such processes may contribute to the oxidation of hypolimnetic materials, providing an expansive anaerobic yet respirable environment for diverse electrogens to consume fermentation products like acetate that could otherwise be consumed by methanogens.

## ACKNOWLEDGEMENTS

We give special thanks to undergraduate researchers Kaela Amundson, Kali Denis, Roger Ort, Zan Abbas, and Mason Polencheck for their numerous contributions to LTER field research, data collection, and work on explorative research fellowship projects. We thank the University of Wisconsin (UW)-Trout Lake Station, the UW Center for Limnology, and the John and Patricia Lane Award program for their invaluable support. We give extra special thanks to Paul Schramm, Gretchen Gerrish, Mike Coakley, Susan Knight, Noah Lottig, Amber Mrnak, Carol Warden, and Pam Fashingbauer for their years of generously collaborative fieldwork advice, support, and important data collection. We thank the U.S. National Science Foundation NTL-LTER site (DEB-0217533, DEB-0822700, DEB-1440297, DEB-2025982) for providing funding of the Microbial Observatory for LTER of Trout Bog and additional bog lakes, as well as by MCB-9977903 and DEB-0702395. We thank the U.S. Department of Energy Joint Genome Institute, a DOE Office of Science User Facility, supported by the Office of Science of the U.S. DOE operated under Contract No. DE-AC02-05CH11231, for sequencing and assembly (CSPs 394 and 2796). This research was also performed in part using the Wisconsin Energy Institute computing cluster, which is supported by the Great Lakes Bioenergy Research Center as part of the U.S. Department of Energy Office of Science. Also, we are thankful for fellowships provided through the department of Bacteriology at UW-Madison and funding from the Wisconsin Alumni Research Foundation.

